# *In vitro* methylation of the U7 snRNP subunits Lsm11 and SmE by the PRMT5/MEP50/pICln methylosome

**DOI:** 10.1101/2023.05.10.540203

**Authors:** Xiao-cui Yang, Anthony Desotell, Min-Han Lin, Andrew S. Paige, Agata Malinowska, Yadong Sun, Wei Shen Aik, Michał Dadlez, Liang Tong, Zbigniew Dominski

## Abstract

U7 snRNP is a multi-subunit endonuclease required for 3’ end processing of metazoan replication-dependent histone pre-mRNAs. In contrast to the spliceosomal snRNPs, U7 snRNP lacks the Sm subunits D1 and D2 and instead contains two related proteins, Lsm10 and Lsm11. The remaining five subunits of the U7 heptameric Sm ring, SmE, F, G, B and D3, are shared with the spliceosomal snRNPs. The pathway that assembles the unique ring of U7 snRNP is unknown. Here, we show that a heterodimer of Lsm10 and Lsm11 tightly interacts with the methylosome, a complex of the arginine methyltransferase PRMT5, MEP50 and pICln known to methylate arginines in the C-terminal regions of the Sm proteins B, D1 and D3 during the spliceosomal Sm ring assembly. Both biochemical and Cryo-EM structural studies demonstrate that the interaction is mediated by PRMT5, which binds and methylates two arginine residues in the N-terminal region of Lsm11. Surprisingly, PRMT5 also methylates an N-terminal arginine in SmE, a subunit that does not undergo this type of modification during the biogenesis of the spliceosomal snRNPs. An intriguing possibility is that the unique methylation pattern of Lsm11 and SmE plays a vital role in the assembly of the U7 snRNP.

## INTRODUCTION

Spliceosomal snRNAs generated by RNA polymerase II (U1, U2, U4, U5, U11, U12 and U4_atac_) contain a conserved 9-nucleotide Sm binding site, AAUUU(U/G)UGG, that nucleates the assembly of a protein ring composed of seven Sm subunits: D1, D2, B, D3, E, F and G (Raker et al. 1999; Will and Luhrmann 2001; Khusial et al. 2005). Each Sm protein occupies a specific position in the ring and contacts an individual nucleotide of the Sm-binding site (Kambach et al. 1999; Khusial et al. 2005; Leung et al. 2011; Li et al. 2016). The Sm proteins exist in the cytoplasm as three preformed heterooligomers, SmD1/D2, SmB/D3 and SmE/F/G (Raker et al. 1996) that are assembled around the Sm-binding site of the spliceosomal snRNAs in a multi-step process mediated by the PRMT5 methylosome complex and the survival of motor neurons (SMN) complex (Battle et al. 2006a; Li et al. 2014; Gruss et al. 2017). Following the assembly of the Sm ring, spliceosomal snRNPs undergo additional maturation steps in the cytoplasm and the nucleus, including binding of specific accessory proteins, to become competent for their role in pre-mRNA splicing (Lührmann et al. 1990; Matera et al. 2007; Wilkinson et al. 2020).

The PRMT5 methylosome consists of PRMT5 (protein arginine methyltransferase 5), MEP50 and pICln (Pu et al. 1999; Friesen et al. 2001b; Meister et al. 2001; Friesen et al. 2002; Gruss et al. 2017). Both biochemical and structural studies indicate that PRMT5 and MEP50 exist as a stable complex of 4 heterodimers, which recruits four pICln molecules through direct interaction with PRMT5 (Antonysamy et al. 2012; Ho et al. 2013; Timm et al. 2018; Mulvaney et al. 2021). PRMT5 (previously termed JBP1), the catalytic component of the complex, belongs to type II protein methyltransferases and generates monomethyl arginines (MMA) as intermediates and symmetric dimethyl arginines (sDMA) as final products (Bedford and Richard 2005). As most protein methyltransferases, PRMT5 has specificity for protein regions rich in glycines and arginines (GR-rich) (Najbauer et al. 1993; Stopa et al. 2015; Musiani et al. 2019). MEP50 (Meister et al. 2001; Friesen et al. 2002), also known as WD45, is a WD40-repeat protein and functions to enhance substrate specificity and catalytic activity of PRMT5 (Burgos et al. 2015). pICln is an adaptor component of the methylosome and delivers the SmD1/D2 and SmB/D3 heterodimers for PRMT5-catalyzed symmetrical arginine dimethylation of the GR-rich C-terminal tails of the Sm subunits D1, B and D3 (Chari et al. 2008; Pesiridis et al. 2009).

The seven Sm proteins, including the PRMT5-modified SmD1, SmB and SmD3, are arranged around the spliceosomal snRNAs by the SMN complex during a multis-step and a highly controlled process. One of these steps is the formation of a ring-shaped 6S intermediate consisting of the Sm proteins D1, D2, E, F and G, and pICln. In the 6S intermediate, pICln temporarily substitutes for the missing SmB/D3 heterodimer and prevents incorporation of illegitimate RNAs (Chari et al. 2008; Grimm et al. 2013). The SMN complex consists of the SMN protein of 30 kDa, Gemins2-8 and Unrip, with the function of only some of them being understood (Battle et al. 2006a; Gruss et al. 2017). The SMN protein is at the center of the complex and acts by directly binding PRMT5- methylated residues (Friesen et al. 2001a; Selenko et al. 2001; Cote and Richard 2005; Tripsianes et al. 2011), hence bringing multiple components of the SMN complex to the vicinity of the Sm proteins. Among these components, Gemin5 recognizes spliceosomal snRNAs as correct assembly targets by simultaneously binding to their monomethylated cap structure and Sm site (Battle et al. 2006b; Lau et al. 2009; Yong et al. 2010; Wahl and Fischer 2016). Gemin2 binds and stabilizes the SmD1/D2/E/F/G pentamer, making extensive contacts with all five subunits (Zhang et al. 2011; Grimm et al. 2013; Yi et al. 2020). This step likely occurs after the departure of pICln from the 6S intermediate and prior to the delivery of a spliceosomal snRNA by Gemin5. The assembly process is completed by the ring closure upon the recruitment of symmetrically dimethylated SmB/D3 heterodimer.

Deficiency of functional SMN protein results in spinal muscular atrophy (SMA), a genetic disorder characterized by selective degeneration of motor neurons and progressive paralysis (Lefebvre et al. 1995; Gubitz et al. 2004; Iannaccone et al. 2004; Pellizzoni 2007; Chari et al. 2009). It is believed that at the molecular level, SMA is caused by inefficient assembly of the spliceosomal snRNPs and aberrant splicing of mRNA precursors important for the development and function of motor neurons (Winkler et al. 2005; Zhang et al. 2008; Lotti et al. 2012). Alternatively, the SMN protein, besides its universal role in snRNP biogenesis, may have a tissue-specific function in motor neurons, explaining selective death of only this group of cells in SMA patients (Monani 2005; Burghes and Beattie 2009; Fallini et al. 2012).

In addition to the spliceosomal snRNPs, animal cells (but not plants or lower eukaryotes) contain the U7 snRNP in which Sm protein D1 and D2 are replaced by the Sm-like proteins Lsm10 and Lsm11, respectively (Schumperli and Pillai 2004; Dominski and Marzluff 2007; Gruss et al. 2017). U7 snRNP is a multi-subunit RNA-guided endonuclease that functions in 3’ end processing of replication-dependent histone pre-mRNAs (Sun et al. 2020; Dominski and Tong 2021), generating mature histone mRNAs that end with a conserved stem-loop structure rather than a poly(A) tail typical of the vast majority of eukaryotic mRNAs (Mandel et al. 2008; Liu and Moore 2021). While Lsm10 is relatively small, resembling in size most Sm proteins (Pillai et al. 2001), Lsm11 has an extended N-terminal region of about 150 amino acids that is essential for the activity of U7 snRNP in 3’ end processing (Pillai et al. 2003; Sun et al. 2020). This region of Lsm11 interacts with FLASH (Yang et al. 2009), facilitating the recruitment of CPSF73, the catalytic subunit of the U7 snRNP (Dominski et al. 2005; Yang et al. 2013; Sun et al. 2020). U7 snRNA, the RNA component of U7 snRNP, consists of ∼60 nucleotides and contains an unusual Sm binding site, AAUUUGUCUAG, that differs from that in the spliceosomal snRNAs and promotes the incorporation of Lsm10 and Lsm11 instead of SmD1 and SmD2 into the U7 snRNP Sm ring (Stefanovic et al. 1995; Schumperli and Pillai 2004).

Initial studies on U7 snRNP suggested that its assembly is at least partially similar to the assembly of the spliceosomal snRNPs, involving both the PRMT5 and SMN proteins (Pillai et al. 2003; Schumperli and Pillai 2004; Azzouz et al. 2005; Li et al. 2014). More recent studies supported this conclusion by showing that pICln recruits the Lsm10/11 heterodimer to the PRMT5/MEP50 heterooctamer (Paknia et al. 2016) and that downregulation of the SMN protein impairs the assembly of U7 snRNP, resulting in a defect in 3’ end processing of histone pre-mRNAs (Tisdale et al. 2013; Tisdale et al. 2022). However, an open question was whether Lsm10 and/or Lsm11 become symmetrically dimethylated by PRMT5 or the interaction of the Lsm10/11 heterodimer with the methylosome plays other than catalytic role(s) in U7 snRNP assembly, contrasting with the assembly of the spliceosomal snRNPs (Azzouz et al. 2005). It was also unclear whether besides SMN, other components of the SMN complex have any role in the assembly of U7 snRNP. *In vivo* studies failed to detect binding of Gemin5 to U7 snRNA (Battle et al. 2006b), suggesting that at least this protein is dispensable for U7 snRNP assembly, being substituted by a counterpart that recognizes the unique Sm-binding site in U7 snRNA and promotes the assembly of the U7-specific ring containing Lsm10 and Lsm11.

Here, we show that the Lsm10/11 heterodimer tightly interacts with both endogenous and recombinant PRMT5 methylosome. Using biochemical and structural approaches, we mapped the binding site of PRMT5 to a short RG cluster within the unique N-terminal region of Lsm11. We also show that two arginine residues in this cluster, but not in other parts of Lsm11 or Lsm10, are methylated *in vitro*. Most surprisingly, within a complex consisting of Lsm10/11 dimer, SmE/F/G heterotrimer and pICln (which we will refer to as the U7 6S intermediate), methylation also occurs at a single arginine residue located near the N-terminus of SmE. Methylation of SmE has never been detected in the context of spliceosomal snRNPs. This residue and the following glycine are conserved in all known vertebrate SmE orthologues, suggesting that symmetric methylation at this site in conjunction with methylation of the N-terminal Lsm11 may provide an important structural determinant that discriminates between the pathways for the assembly of the U7 snRNP and the spliceosomal snRNPs.

## RESULTS

### Recombinant Lsm10/11 heterodimer interacts with endogenous PRMT5 methylosome from mammalian extracts

Our recent biochemical and structural studies with recombinant U7 snRNP uncovered key interactions within this muti-subunit endonuclease and demonstrated how it cleaves histone pre-mRNAs (Sun et al. 2020; Yang et al. 2020). We next focused on the biogenesis of the U7 snRNP, including the poorly understood process of assembling U7-specific Sm ring on U7 snRNA. To facilitate this study, we generated rabbit antibodies against baculovirus-expressed heterodimer of Lsm10-MBP fusion protein and His-tagged Lsm11. The resultant serum from immunized rabbits (α10/11) precipitated from the cytoplasm of HeLa cells very small amounts of endogenous Lsm10 and Lsm11 that could not be visualized by silver staining (Fig. 1A, lane 1), but were readily detectable by Western blotting (data not shown). Interestingly, when the same HeLa cytoplasm was supplemented with Lsm10-MBP/Lsm11 heterodimer, the α10/11 serum in addition to the two recombinant proteins precipitated three endogenous proteins strongly stained with silver (Fig. 1A, lane 2). They were identified by mass spectrometry and Western blotting as subunits of the PRMT5 methylosome: PRMT5 methyltransferase of ∼70 kDa (Stopa et al. 2015), MEP50 of ∼50 kDa (Friesen et al. 2002) and ∼35 kDa pICln (Pu et al. 1999; Meister et al. 2001). PRMT5 and MEP50 form a heterooctamer consisting of 4 copies of each protein (Antonysamy et al. 2012; Ho et al. 2013; Timm et al. 2018), and pICln binds stoichiometrically to PRMT5 (Friesen et al. 2001b; Pesiridis et al. 2009; Guderian et al. 2011; Mulvaney et al. 2021), explaining the abundance of the three methylosome subunits relative to the small amounts of Lsm10 and Lsm11 used in the assay.

**Fig. 1.**
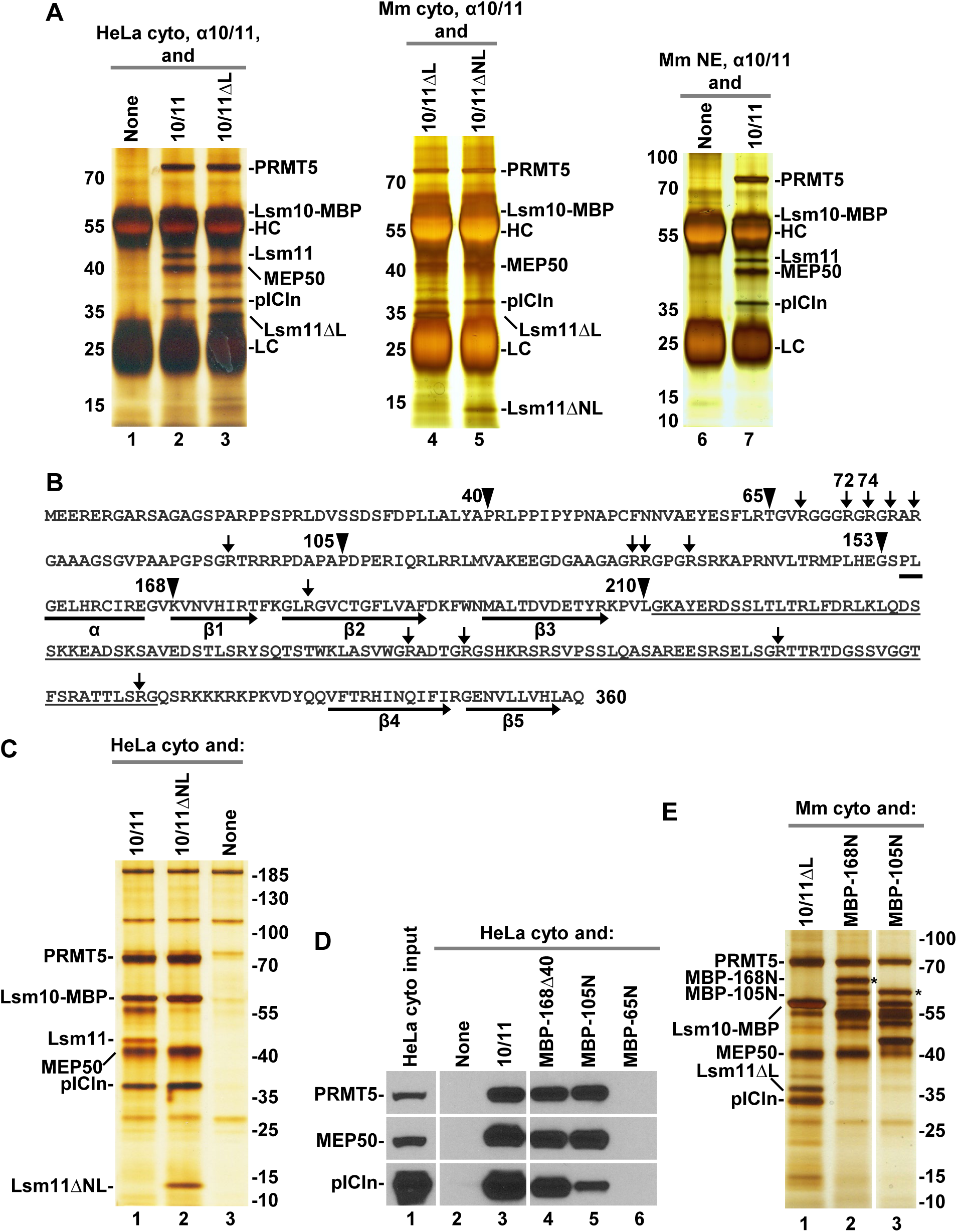
Lsm11 interacts with the PRMT5 methylosome. **A.** Immunoprecipitation of HeLa (lanes 1-3) or mouse proteins (lanes 4-7) bound to recombinant heterodimers of Lsm10 with either wild type Lsm11 (labelled as 10/11), Lsm11 lacking the internal loop (10/11ΔL), or Lsm11 lacking both the N-terminal fragment and to loop (10/11ΔNL). The heterodimers were incubated with indicated extracts and precipitated with the α10/11 rabbit serum. Immunocomplexes were collected on protein A beads, resolved by SDS-PAGE and protein bands stained with silver. Proteins precipitated by the same serum from HeLa or mouse extracts lacking recombinant Lsm10/11 heterodimer are shown in lanes 1 and 6. HC and LC denote heavy and light chains of immunoglobulins, respectively. **B.** Sequence of human Lsm11 (amino acid 1-360). The boundaries of the deleted regions are indicated with arrowheads. Arginine residues neighboring glycines potentially targeted for methylation by PRMT5 are indicated with arrows. Structural elements within the Sm fold are indicated with a horizontal bar (α-helix) or thick arrows (β-stands). The portion of the internal loop between β-stands 3 and 4 that was deleted in Lsm11ΔL and Lsm11ΔNL is underlined. **C-E.** Purification of PRMT5 methylosome on amylose beads via Maltose Binding protein (MBP) tag attached to Lsm10 or to the N-terminal fragments of Lsm11. HeLa or mouse cytoplasmic extracts, as indicated, were incubated with variants of Lsm10/11 heterodimers or N-terminal fragments of Lsm11 fused to MBP and proteins collected on amylose beads were analyzed by silver staining (panels C and E) or Western blotting, using specific antibodies (panel D). In panel E, asterisks indicate full length N-terminal Lsm11 fragments fused to MBP.

Human Lsm11 with its 360-amino acids is the largest member of the Sm/Lsm family and contains an extended N-terminal region of 154 residues that plays a critical role in the function of U7 snRNP in 3’ end processing (Fig. 1B). Within this region, amino acids ∼20-50 bind FLASH (Yang et al. 2009) to form a platform for recruiting the CPSF73 endonuclease and two other component of the cleavage and polyadenylation machinery: CPSF100 and symplekin (Yang et al. 2013; Aik et al. 2017). Between amino acids 68-79, Lsm11 contains a GR cluster that may serve as a primary methylation target for PRMT5. Amino acids 155-360 of Lsm11 fold into a canonical Sm domain consisting of an N-terminal α helix (amino acids 155-165) and five β strands (Sun et al. 2020). The β3 and β4 strands are separated by a large loop (amino acids 207-337) that is unique to Lsm11 and contains four separate GR or RG dipeptide motifs (Fig. 1B).

To identify regions of Lsm11 that bind the PRMT5 methylosome, we expressed Lsm10 fused to MBP in complex with Lsm11 lacking the internal loop (amino acids 211- 322, Lsm11ΔL), or Lsm11 lacking both the loop and the entire N-terminal region encompassing amino acids 1-154 (Lsm11ΔNL) (Fig. S1). We also bacterially expressed four N-terminal fragments of Lsm11 fused to MBP (Fig. S1). These fragments lack the Sm fold region of Lsm11 that interacts with Lsm10 and were expressed alone.

Deletion of the internal loop from Lsm11 had no effect on the interaction of the Lsm10/11 heterodimer with endogenous PRMT5 methylosome. Compared to the wild-type dimer containing full length Lsm11, approximately the same amounts of PRMT5, MEP50 and pICln were co-precipitated together with Lsm10/11ΔL dimer by α10/11 serum (Fig. 1A, lane 3). Thus, the extensive loop despite containing several GR/RG dipeptide motifs does not bind the methylosome. Readily detectable binding of the methylosome to the Lsm10/11ΔL dimer was also observed in a cytoplasmic extract from mouse myeloma cells (Fig. 1A, lane 4) and was not abolished by additionally deleting the entire N-terminal region of Lsm11 (Fig. 1A, lane 5). Finally, binding of PRMT5, MEP50 and pICln to Lsm10/11 heterodimers was observed in a mouse myeloma nuclear extract (Fig. 1A, lane 7), indicating that the interaction can occur in both the cytoplasmic and nuclear fractions from different mammalian cell lines.

To further analyze the interaction between the Lsm10/11 dimer and the PRMT5 methylosome, we tested a different approach by directly collecting proteins bound to the dimer on amylose beads via the MBP tag attached to Lsm10 rather than using immunoprecipitation with α10/11 serum. Following a brief incubation of the wild type or Lsm11ΔNL heterodimer with a HeLa cytoplasmic extract, the bound proteins were collected on amylose beads and visualized by silver staining following their separation by SDS-PAGE. Again, both dimers, one containing full length Lsm11 and the other lacking the N-terminus and the internal loop co-purified with large amounts of the PRMT5 methylosome (Fig. 1C, lanes 1 and 2, respectively). In the absence of Lsm10/11 dimers only a background of several protein bands was detected but no trace of the three methylosome components (Fig. 1C, lane 3).

We used the same approach with various N-terminal fragments of Lsm11 tagged at the N-terminus with MBP: MBP-Δ40 (Lsm11 amino acids 41-169), MBP-105N (Lsm11 amino acids 1-105) and MBP-65N (Lsm11 amino acids 1-65) (Fig. S1). As determined by silver staining (not shown) and Western blotting, MBP-Δ40 and MBP-105N clearly interacted with PRMT5 and MEP50 (Fig. 1D, lanes 4-5), but compared to the Lsm10/11 dimer, co-precipitated significantly less of pICln (Fig. 1D, lane 3), consistent with this adaptor protein contacting Lsm10 (see below). No trace of the methylosome was detected on amylose beads in the absence of recombinant proteins or in the presence of the first 65 amino acids of Lsm11 fused to MBP (MBP-65N) (Fig. 1D, lanes 2 and 6, respectively).

We repeated this experiment using a cytoplasmic extract from mouse myeloma cells rather than from Hela cells and an additional Lsm11 protein encompassing all 168 N-terminal amino acids of the protein. Again, as visualized by silver staining, the heterodimer (in this case Lsm10/11ΔL) was very efficient in binding pICln (Fig. 1E, lane 1). MBP- 168N and MBP-105N pulled down only a trace of this subunit while binding efficiently the two remaining proteins of the methylosome, PRMT5 and MEP50 (Fig. 1E, lanes 2 and 3). Altogether, these results indicate that Lsm10/11 contains at least two binding sites for the methylosome. The first site is located within the region delineated by amino acids 65 and 105. This region contains an RG cluster (amino acids 58-79) that likely interacts with PRMT5 (Friesen et al. 2001b). The second site likely corresponds to the Sm fold of Lsm10 that tightly interacts with pICln, resembling the interaction of this subunit with SmD1 (Chari et al. 2008; Grimm et al. 2013; Paknia et al. 2016).

### Binding of recombinant PRMT5 methylosome to Lsm10/11 heterodimer and methylation of Lsm11

To determine whether the interaction between PRMT5 methylosome and Lsm10/11 can be reconstituted from recombinant components, we co-expressed PRMT5 and MEP50 in baculovirus system, and purified pICln as a separate protein from bacteria. The PRMT5/MEP50 complex was mixed with the Lsm10/11 heterodimer in molar ration 1:1 either in the absence or in the presence of pICln, and the amount of the methylosome components immobilized on amylose beads via the MBP tag attached to Lsm10 was assessed by SDS-PAGE and silver staining. As seen in Fig. 2A, both PRMT5 and MEP50 were readily detected and their amount increased in the presence of pICln, consistent with its role as the adaptor subunit for recruiting the Lsm10/11 substrate. We conclude that the interaction between Lsm10/11 heterodimer and the methylosome can be recapitulated using recombinant components.

**Fig. 2.**
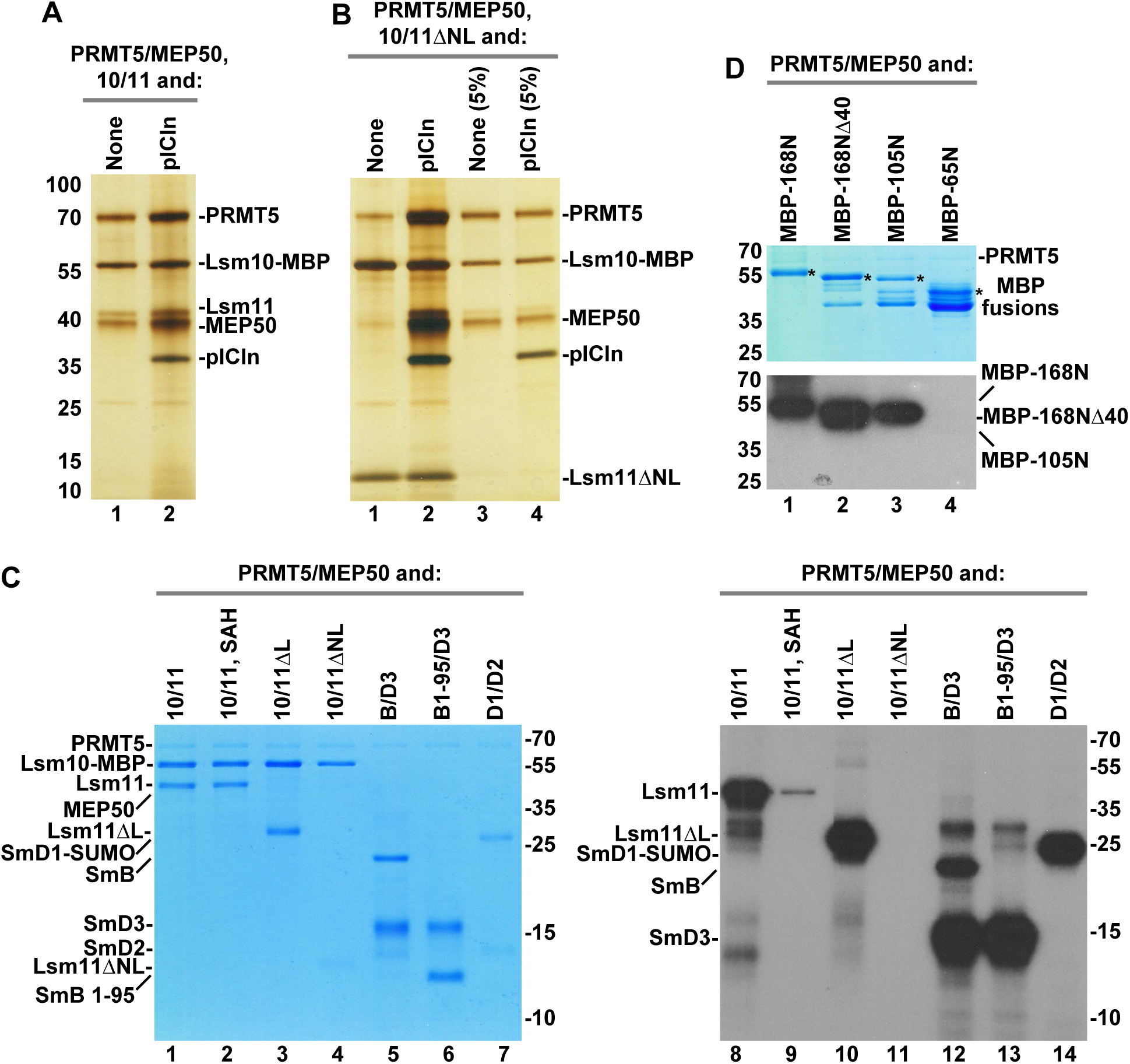
Binding and methylation activities of recombinant PRMT5 methylosome. **A.** PRMT5/MEP50 complex without (lane 1) or with pICln (lane 2) bound to Lsm10/11 heterodimer was purified on amylose beads via MBP attached to Lsm10 and analyzed by silver staining. **B.** Lsm10/11ΔNL heterodimer was tested for the ability to stably bind PRMT5/MEP50 complex either in the absence (lane 1) or in the presence of pICln (lane 2). Proteins purified on amylose beads via MBP attached to Lsm10 were visualized by silver staining. Lanes 3-4 represent 5% input of proteins used in the binding assay. **C.** Various Sm or Lsm heterodimers were incubated overnight in solution with recombinant complex of PRMT5 and MEP50 in the presence of ^3^H SAM. Proteins were resolved by SDS-PAGE and visualized by staining with Coomassie Blue (lanes 1-7). Dried gel was used for fluorography to detect proteins labeled with radioactive methyl group (lanes 8- 14). **D.** Various N-terminal fragments of Lsm11 fused at the N-terminus with MBP were tested for their ability to undergo methylation, as described in panel C. Coomassie Blue stained gel and fluorogram are shown at the top and bottom, respectively. Asterisks in the top panel indicate full length N-terminal Lsm11 fragments fused to MBP. Lower bands correspond to proteolytic fragments.

Our studies with cytoplasmic extracts suggested that binding of endogenous methylosome to the Lsm10/11 heterodimer with the deletion of both the internal loop and the N-terminal region (Lsm10/11ΔNL), hence lacking the RG cluster between residues 58- 79, likely depends on pICln, which is predicted to bind the Sm fold of Lsm10 (Paknia et al. 2016). With the availability of recombinant PRMT5/MEP50 heterodimer and pICln as two separate entities, we directly tested this interpretation by either adding or omitting pICln in the binding assay with the Lsm10/11ΔNL heterodimer. Only background amounts of PRMT5 and MEP50 bound to Lsm10/11ΔNL heterodimer when pICln was excluded and the interaction was greatly stabilized in its presence (Fig. 2B, lanes 1 and 2, respectively). Thus, as with endogenous methylosome, recombinant methylosome recognizes two independent binding sites in the Lsm10/11 heterodimer, one likely being the RG cluster (amino acids 58-79) that interacts with PRMT5, and the other being the Sm fold of Lsm10 that interacts with pICln. These two contacts collectively contribute to the strong interaction between the methylosome and the Lsm10/11 heterodimer.

The strong association of the PRMT5 methylosome with the Lsm10/11 heterodimer and with the N-terminal region of Lsm11 containing a GR cluster raised the possibility that either Lsm11 alone or both subunits of the heterodimer are methylated. This would be consistent with the catalytic activity of PRMT5 on the RG-rich C-terminal tails of SmD1, SmB and SmD3. To test this possibility directly, we carried out an *in vitro* methylation assay using all recombinant components and ^3^H-labeled SAM as a source of radioactively labelled methyl group. Initially, the methylation reaction was carried out only with PRMT5 and MEP50, in the absence of pICln. We tested methylation of the three variants of Lsm10/11 heterodimers, in which Lsm11 was either full length or lacked the N-terminal region and/or the internal loop, as described above (Fig. S1). As a control, we used spliceosomal heterodimers SmD1/D2 and SmB/D3 that are known substrates for *in vitro* symmetric arginine dimethylation by the PRMT5 methylosome (Friesen et al. 2001b; Meister et al. 2001; Friesen et al. 2002). As an additional control, we used a variant of SmB/D3 heterodimer (SmB1-95/D3) in which the SmB subunit encompassed amino acids 1-95, lacking the entire unstructured GR-rich C-terminal tail.

Following overnight incubation, all samples were resolved by SDS-PAGE and stained with Coomassie Blue to visualize proteins (Fig. 2C, lanes 1-7) and subsequently used fluorography to determine their methylation status (Fig. 2C, lanes 8-14). Incubation of PRMT5 and MEP50 with Lsm10/11 and Lsm10/11ΔL heterodimers resulted in methylation of both full length Lsm11 and Lsm11ΔL, but not Lsm10 (Fig. 2C, lanes 8 and 10). As expected, radioactive labelling was almost completely blocked by excess of the competitive methylation inhibitor SAH (Fig. 2C, lane 9) (Meister et al. 2001). Thus, Lsm11, but not Lsm10, is a substrate for *in vitro* methylation by PRMT5, with methylation occurring somewhere outside the internal loop. Methylation was entirely abolished in Lsm10/11ΔNL heterodimer (Fig. 2C, lane 11), demonstrating that the methylation site is located within the deleted N-terminal portion of Lsm11. The spliceosomal Sm proteins, D1, B and D3 were methylated as expected, with a deletion of the C-terminal tail abolishing methylation of SmB (Fig. 2C, lanes 12-14).

To map the methylation site more precisely, we tested N-terminal fragments of Lsm11 fused at the N-terminus to MBP (Fig. S1). Again, the reaction was carried in in the presence of ^3^H SAM and limiting amounts of recombinant PRMT5 and MEP50. Of the four N-terminal fragments of Lsm11 fused to MBP, 168N, 168NΔ40 and 105N, became radioactively labeled (Fig. 2D, lane 1-3), whereas the shortest 65N fragment was not methylated (Fig. 2D, lane 4). In contrast to the three other fragments, MBP-65N does not bind the methylosome and lacks the GR cluster, strengthening the notion that methylosome binding site and the methylation site are located between amino acids 68-79 of Lsm11.

### Structure of the methylosome-Lsm10/11 complex

To determine how PRMT5 interacts with Lsm11, we mixed all three methylosome components with the Lsm10/11 heterodimer and purified their complex by gel filtration chromatography for cryo-EM studies, which yielded a structure of the complex at 2.86 Å resolution (Fig. 3A-D, Table 1). Clear EM density was observed for PRMT5 and MEP50, and for only a few residues of the PRMT5 binding motif (PBM) in pICln (Mulvaney et al. 2021) (Fig. 3A). The remaining parts of pICln, most of Lsm11 and the entire Lsm10 were disordered, providing no structural information about the interaction between the Lsm10/Lsm11 heterodimer and pICln. PRMT5 and MEP50 form a heterooctamer of four tightly bound heterodimers (Fig. 3A), consistent with previous structural studies of the methylosome by X-ray crystallography and cryo-EM (Antonysamy et al. 2012; Ho et al. 2013; Timm et al. 2018).

**Fig. 3.**
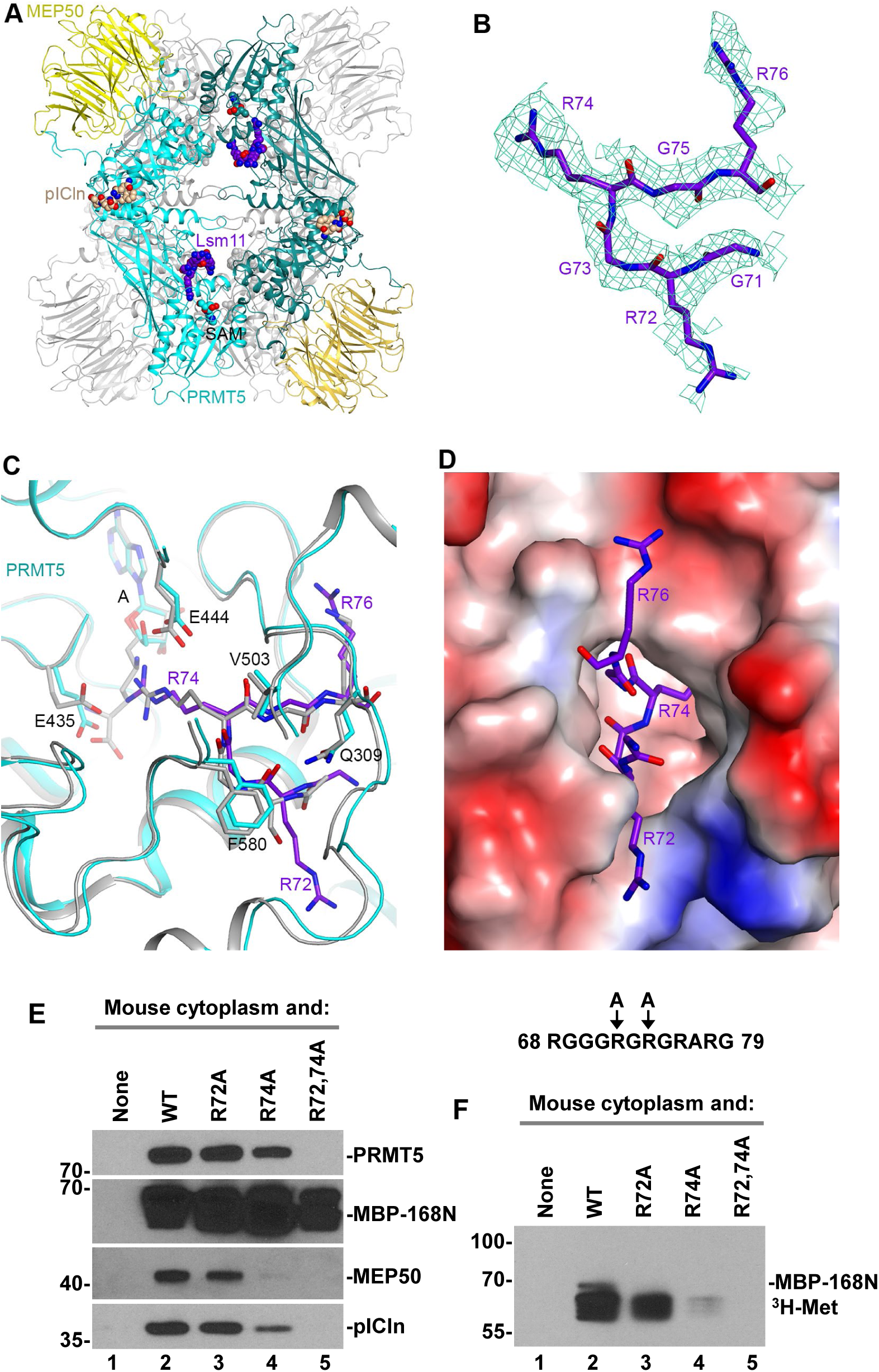
Structure of human methylosome-Lsm10/11 complex. **A.** Schematic drawing of the human methylosome-Lsm10/11 complex. The molecules in the top layer are shown in color, and those in the bottom layer are in gray. Peptide segments from pICln and Lsm11, as well as the adenosine portion of the SAM cofactor, are shown in sphere model. **B.** Cryo-EM density for the Lsm11 peptide in the active site of PRMT5. The fitted atomic model is shown in stick model. **C.** Detailed interactions between the Lsm11 peptide and the active site of PRMT5. The structure of histone H4 peptide bound to the methylosome is shown in overlay (in gray). **D.** Electrostatic surface of the active site region of PRMT5. The Lsm11 peptide is shown in stick model. The structure figures were produced with PyMOL (www.pymol.org). **E and F.** WT Lsm11 encompassing the first 168 amino acids N-terminally fused to MBP (MBP-168N) and indicated Lsm11 mutants were incubated with a mouse cytoplasmic extract. Proteins bound to Lsm11 were immobilized on amylose beads via MBP. The beads were split evenly into two halves and analyzed by Western blotting (panel E) for the presence of methylosome components or resuspended in a buffer containing ^3^H SAM to test methylation of Lsm11, as detected using fluorography (panel F).

We observed EM density consistent with a peptide segment in the active site of PRMT5. After examining many possibilities, residues 71-GRGRGR-76 in the N-terminal extension of Lsm11 were chosen as the most likely interpretation of this density (Fig. 3B). The density exists in the active sites of all four PRMT5 molecules, although the quality of the density varied among them. The side chain of Arg74 is inserted deep into the active site and would be located next to the cofactor SAM (Fig. 3C and D), arguing that it is poised for methylation. Since the density for the side chain of Arg72 supporting this model is relatively weak, we cannot exclude the possibility that the active site of PRMT5 accommodates the overlapping 69-GGGRGR-74 Lsm11 peptide. In this alternative model, methylation would occur at Arg72, a residue that is well conserved among vertebrate Lsm11 orthologues. Plausibly, the observed EM density is derived from a mixture of the two different peptides bound in the active site of different PRMT5 molecules.

The six-residue peptide of Lsm11 bound to the active site of PRMT5 assumes the conformation of a type II reverse turn, with the Arg residue ready for methylation located at the third position of the turn between two glycines in the primary sequence (Fig. 3C). Consistently, there is no indication of EM density for a Cβ atom on either side of the methylated Arg (Fig. 1B). In fact, a Cβ atom on the residue before the methylated Arg would clash with the side chain of Phe580 of PRMT5 (Fig. 3C). For the residue that follows the methylated Arg, a Cβ atom would be located ∼3 Å from the side chains of Gln309 and Val503, and a larger side chain here would clash with PRMT5. Therefore, the current structure suggests that a GRG motif may be preferred for the methylation site, and a GRA motif could also be accommodated.

The binding mode of the Lsm11 peptide resembles that of the SGRGK histone H4 peptide (Antonysamy et al. 2012), in which the methylated arginine is also flanked by two glycines. The overall structures of the two complexes are similar, with rms distance of 0.64 Å for 609 equivalent Cα atoms of PRMT5 and 0.36 Å for 295 equivalent Cα atoms of MEP50. For the cofactor SAM (or SAH), EM density for only adenosine is observed in the four active sites of PRMT5, and the rest of the cofactor (the Met residue) is disordered (Fig. 3C). Possibly related to this disordering, the side chain of Glu435 does not interact with Arg74 of Lsm11 and instead assumes a different conformation that would clash with the main chain of the Met residue of SAM (Fig. 3C).

To determine which of the arginines in the RG cluster of Lsm11 is methylated, we generated three mutants of the N-terminal Lsm11 region (amino acids 1-168) fused N-terminally to MBP (MBP-168N): R72A, R74A and double mutant R72,74A. In these mutants, either one or two arginines predicted by cryo-EM to undergo methylation were replaced with alanines. Both the wild type MBP-168N and three mutant proteins were incubated with a cytoplasmic extract from mouse myeloma cells and proteins immobilized on amylose beads via the MBP were visualized by silver staining (not shown) and Western blotting (Fig. 2E). As expected, no trace of PRMT5, MEP50 and pICln was recovered from the beads in the absence of MBP-168N, and the wild type protein readily bound all three methylosome subunits (Fig. 3E, lanes 1 and 2). The R72A mutant was almost as efficient in binding the methylosome, whereas the R74A mutant bound weakly, as illustrated by significantly reduced amounts of MEP50 and pICln (Fig. 3E, lanes 3 and 4). Importantly, the R72,74A double mutant failed to interact with the methylosome (Fig. 3E, lane 5).

We also tested the ability of each sample to methylate Lsm11. The beads were resuspended in a buffer containing ^3^H SAM and incubated overnight. As expected, results of this *in vitro* methylation reaction mirrored the results of binding assay, with the strongest radioactive signal observed for the wild type MBP-168N. The R72A mutant was methylated with a lower efficiency, whereas the radioactive signal for R74A and R72,74A mutants was very weak and undetectable, respectively (Fig. 3F). Based on the cryo-EM reconstruction, binding assay, and *in vitro* methylation, we conclude that arginine in position 74 of Lsm11 is the primary binding and methylation target for the methylosome. PRMT5 also binds and methylates arginine 72, but with lower efficiency than that observed for arginine 74.

### Methylation of SmE

*In vivo*, SmD1/D2 heterodimer is believed to rapidly associate with pICln, which recruits the heterodimer to the methylosome for methylation of SmD1. The heterodimer subsequently associates with a trimer of SmE, SmF and SmG, forming a stable 6S intermediate complex consisting of Sm proteins D1/D2/E/F/G and pICln (Chari et al. 2008; Grimm et al. 2013; Paknia et al. 2016). We tested whether methylation of Lsm11 can be affected by pICln. We also tested methylation of Lsm11 in the presence of both pICln and the SmE/F/G heterotrimer that together may form a complex equivalent to the spliceosomal-type 6S complex (Neuenkirchen et al. 2015; Paknia et al. 2016).

When the methylation reaction was carried out in the presence of either pICln, which binds Lsm10/11 (Paknia et al. 2016) or SmE/F/G heterotrimer, which based on the behavior of SmD1/D2 is unlikely to form a stable complex with a heterodimer of Lsm10/11 (Chari et al. 2008; Zhang et al. 2011; Grimm et al. 2013), methylation of Lsm11 was virtually unaffected (Fig. 4A, bottom panel, lanes 1-3). Strikingly, when both pICln and the SmE/F/G heterotrimer were included into the reaction, methylation of an additional protein migrating between 10 and 15 kDa size markers was detected (Fig. 4A, bottom panel, lane 4). Alignment of the Coomassie Blue stained gel with the fluorogram tentatively identified the methylated protein as SmE, which migrates at the top of the SmE/F/G triplet (Fig. 4A, top panel, lane 4).

**Fig. 4.**
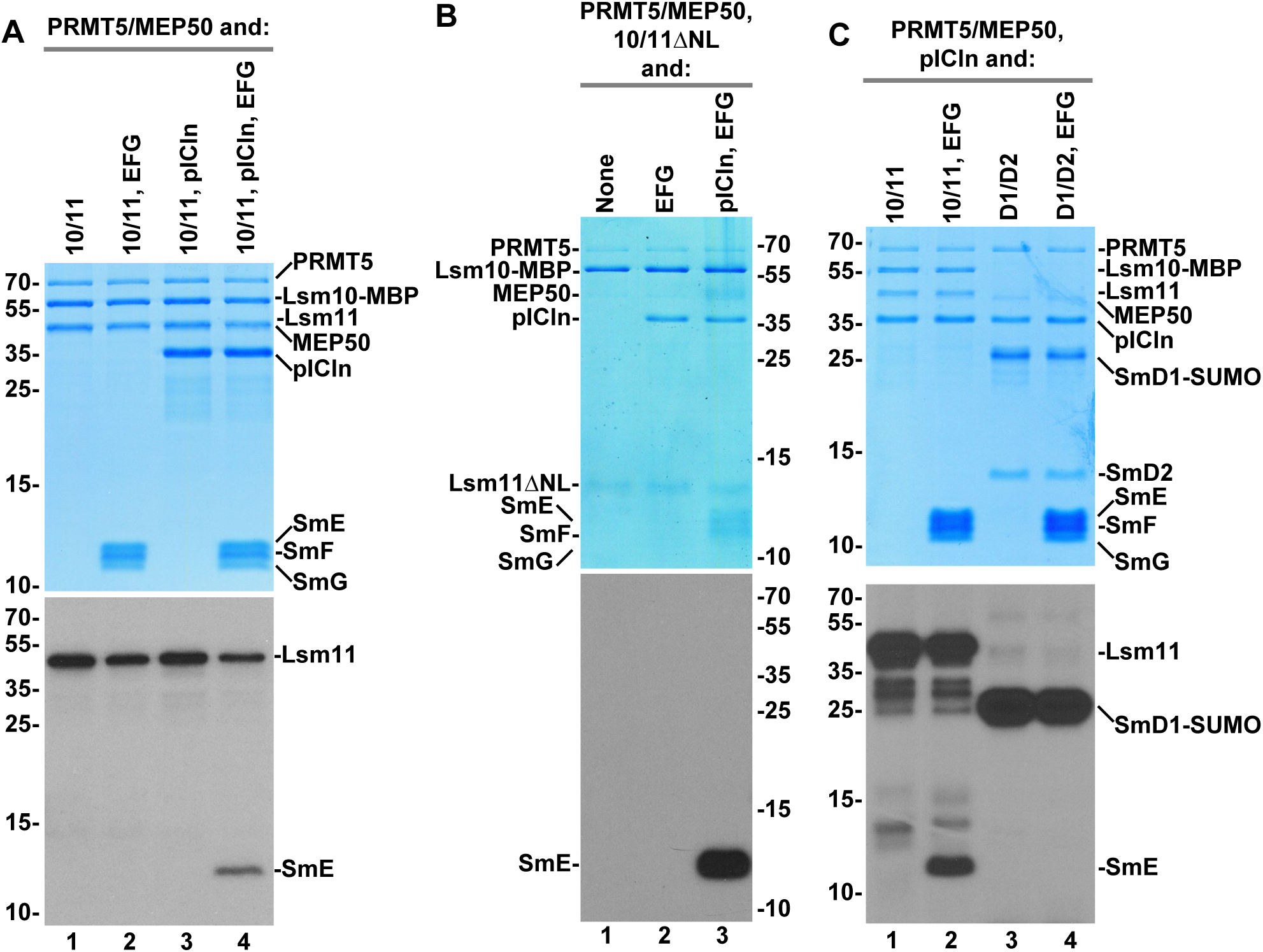
Methylation of SmE by recombinant methylosome. A-C. Recombinant PRMT5/MEP50 complex was incubated with indicated Sm or Lsm heterodimers and ^3^H SAM either in the absence or in the presence of pICln and SmE/F/G heterotrimer, as indicated. Protein methylation was detected, as described in the legend for Fig. 2. Coomassie Blue stained gels and fluorograms are shown at the top and bottom, respectively.

The most obvious difference between the SmD1/D2 and Lsm10/11 heterodimers is the presence of the long N-terminal region in Lsm11, which does not exist in SmD2, the spliceosomal counterpart of Lsm11. To determine whether this Lsm11 region is important for methylation of SmE, we carried out *in vitro* methylation assay using Lsm10/11ΔNL heterodimer, in which Lsm11 lacks both the N-terminal extension and the internal loop within its Sm fold (Fig. S1). As expected, due to the deletion of RG-rich region between amino acids 68-79, no methylation of Lsm11 was observed (Fig. 4B, lanes 1-3). Yet, methylation of SmE in the presence of SmE/F/G and pICln was even more pronounced, being detected as a strong signal after a relatively short exposure. Thus, the N-terminal region of Lsm11 may play an inhibitory role in methylation of SmE, perhaps acting as a competitor for binding PRMT5.

We tested whether methylation of SmE can occur if the Lsm10/11 heterodimer is replaced with the spliceosomal SmD1/D2 heterodimer. Only SmD1 (fused to SUMO) was methylated, as expected, and no radioactive signal was detected for SmE or any other protein of the complex (Fig. 4C, lane 4). In a parallel reaction carried out with Lsm10/11, pICln and SmE/F/G, SmE was readily methylated (Fig. 4C, lane 2). This result is consistent with previous studies demonstrating that only SmD1 becomes methylated by the PRMT5 methylosome in the 6S complex consisting of SmD1/D2 heterodimer, SmE/F/G heterotrimer and pICln (Neuenkirchen et al. 2015). Thus, methylation of SmE appears to be an event specifically linked to Lsm10/11 heterodimer and may play a role in the assembly of U7 snRNP *in vivo*.

### SmE is methylated by endogenous methylosome

We next tested whether methylation of SmE can be carried out by endogenous methylosome. Heterodimers of Lsm10 with either full length or ΔNL Lsm11 were incubated with a mouse cytoplasmic extract either in the absence or in the presence of the SmE/F/G heterotrimer and bound complexes collected on amylose beads via the MBP tag attached to Lsm10, as described above. Following exhaustive washes, the beads were incubated overnight in a buffer containing radioactive SAM and the bound material was separated by SDS-PAGE, stained with Coomassie Blue (Fig. 5A, top) and subsequently analyzed by fluorography (Fig. 5A, bottom). Each recombinant Lsm10/11 heterodimer bound a number of proteins, some of which were identified as the components of endogenous methylosome (Fig. 5A, top panel, lanes 1-4, Fig. S2), consistent with the data shown in Fig. 1C. Importantly, endogenous methylosome in the presence of the SmE/F/G heterotrimer methylated SmE (Fig. 5A, bottom panel, lanes 2 and 4), with methylation of this subunit typically exceeding the level observed for full-length Lsm11 (Fig. 5A, bottom panel, lane 2, Fig. S2, lane 6, and Fig. S3, lane 3), supporting the conclusion that SmE is a legitimate PRMT5 methylation target. Lsm11ΔNL that lacks the N-terminal RG cluster failed to undergo any detectable methylation (Fig. 5A, bottom panel, lanes 3 and 4). Only background proteins and no components of the methylosome were purified from HeLa (Fig. S2) and mouse (Fig. S3) cytoplasmic extracts in the absence of recombinant Lsm10/11 heterodimer. As expected, no protein methylation was detected in these samples.

**Fig. 5.**
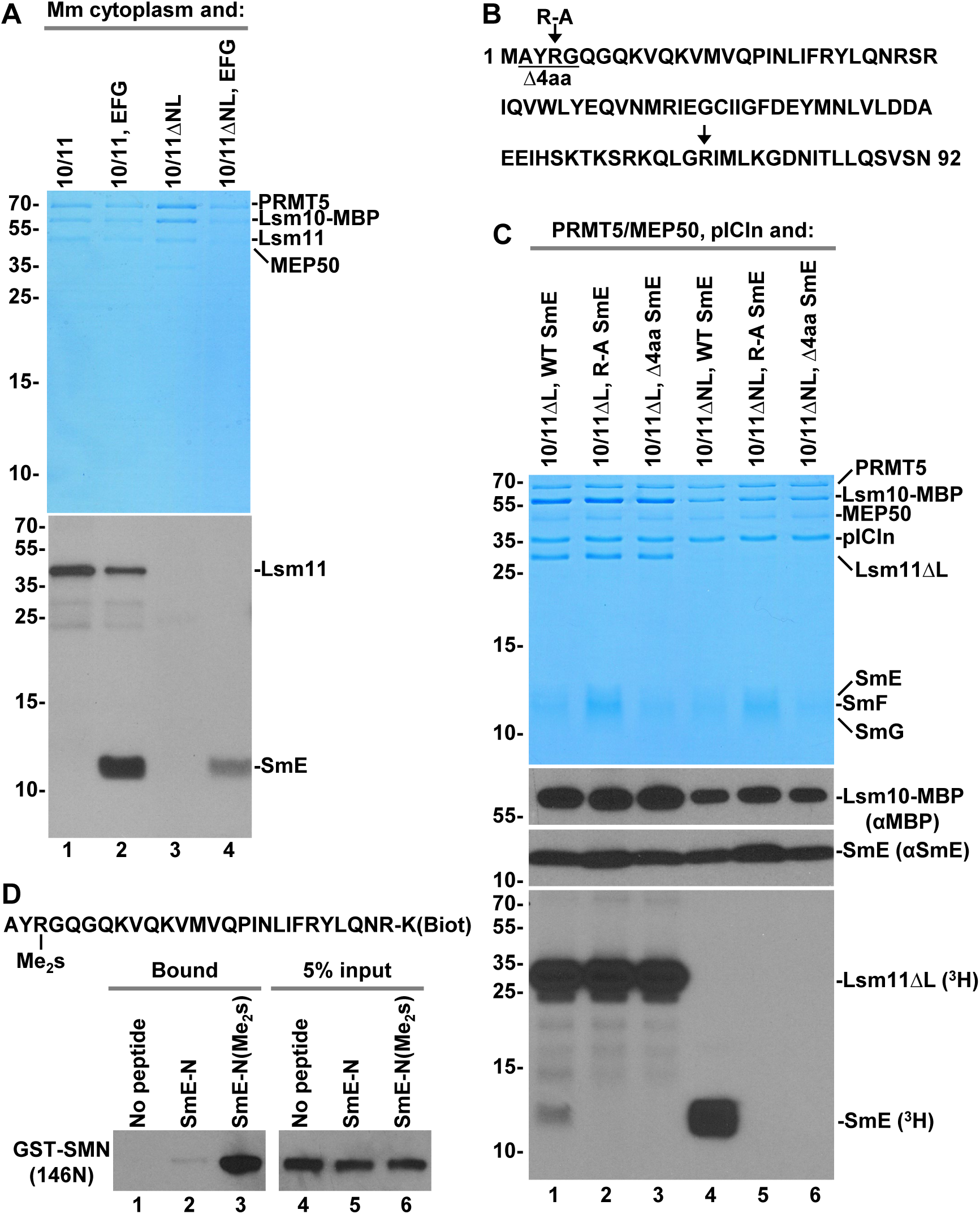
*In vitro* methylation of SmE by endogenous methylosome and identification of the SmE methylation site. **A.** Recombinant Lsm10/11 (lanes 1 and 2) and Lsm10/11ΔNL (lanes 3 and 4) heterodimers were incubated either in the absence or in the presence of the SmE/F/G heterotrimer with a mouse cytoplasmic extract and purified together with bound endogenous methylosome on amylose beads via MBP attached to Lsm10. The immobilized proteins were tested directly on the beads in a buffer containing ^3^H SAM for the ability to methylate Lsm11 and SmE. Following overnight methylation, proteins in each sample were stained with Coomassie Blue (top panel) and their methylation status analyzed by fluorography (bottom panel). **B.** Sequence of human SmE (amino acids 1-92) and mutations made near the N-terminus. The arrows indicate each of the two arginines in SmE neighboring a glycine. **C.** Recombinant methylosome complex consisting of PRMT5, MEP50 and pICln was incubated with Lsm10/11ΔNL heterodimer in the presence of ^3^H SAM and SmE/F/G heterotrimer containing WT, R-A or Δ4aa variants of SmE. Proteins used in the assay were resolved by SDS-PAGE and visualized by staining with Coomassie Blue (top panel) and their methylation status analyzed by fluorography (bottom panel). Lsm10-MBP and SmE were additionally detected by Western blotting using αMBP and αSmE antibodies, respectively (middle panels). **D.** Two C-terminally biotinylated peptides encompassing the first 27 amino acids of SmE and either lacking or containing symmetric dimethyl group at Arg4 were incubated with GST-tagged N-terminal region of SMN (amino acids 1-146) containing the Tudor domain. Proteins immobilized on streptavidin beads were resolved by SDS-PAGE and probed using anti-GST antibody. Lane 1 represents the background observed in the absence of any biotinylated peptide. The input (5%) for all binding reactions is shown in lanes 4-6.

### Mutational analysis identifies the site of methylation in SmE

To prove that Lsm10/11 heterodimer promotes methylation of SmE and to identify the site of methylation, we used mutational analysis. The sequence of SmE contains two arginines directly neighboring a glycine residue: an RG dipeptide located near the N-terminus (amino acids 4-5) and a GR dipeptide (amino acids 75-76) located in the β-strand 4 of the Sm fold (Fig. 5B). The N-terminal arginine is located within the region that is likely unstructured and this residue was considered a primary candidate for methylation in SmE. We substituted this arginine with alanine (R-A) or deleted four N-terminal amino acids including the RG dipeptide (Δ4aa) and tested a SmE/F/G heterotrimer containing the resultant SmE mutants for the ability to undergo methylation by recombinant PRMT5 methylosome. Importantly, both mutations tested in the context of heterodimers containing ΔL Lsm11 (Fig. 5C, bottom panel, lanes 1-3) or ΔNL Lsm11 (Fig. 5C, bottom panel, lanes 4-6) abolished methylation of SmE. The effect of these two mutations was particularly striking when tested in the presence of Lsm10/11ΔNL, which promotes very efficient methylation of wild type SmE (Fig. 5C, bottom panel, compare lane 4 with lanes 5-6).

Since the three subunits of the SmE/F/G often migrate as a cluster of weakly separated bands in 15% SDS/polyacrylamide gels and are difficult to identify as individual proteins, we used Western blotting and anti-SmE antibody to confirm the presence of SmE in the samples that failed to show SmE methylation. We also used anti-MBP antibody to confirm the identity of the Lsm10 subunit visualized by Coomassie Blue staining (Fig. 5C, top panel). Bands corresponding to SmE and Lsm10-MBP were detected in all six samples (Fig. 5C, middle panels). Based on these experiments, we conclude that the arginine in position 4 of SmE is the only residue of the SmE/F/G heterotrimer that becomes methylated by PRMT5/MEP50 in the presence of pICln and Lsm10/11 dimer.

The methylation site in SmE in both its location near the N-terminus and the presence of only one flanking glycine sharply contrasts with the methylation sites in Sm protein D1, B and D3, which are located within C-terminal GR-rich regions. Symmetrically dimethylated sites in these Sm subunits are bound by the SMN Tudor domain, facilitating the assembly of the Sm ring on spliceosomal snRNAs (Friesen et al. 2001a; Selenko et al. 2001; Cote and Richard 2005; Tripsianes et al. 2011). We tested whether the unusual methylation site in SmE is recognized by the Tudor domain of SMN. We ordered chemical synthesis of two peptides containing biotin at the C-terminus and encompassing the first 27 amino acids of SmE (Fig. 5D). Of the two peptides, one was unmodified and the other contained symmetric dimethyl group at Arg4. A binding assay was carried out in the presence of each peptide and GST-tagged SMN fragment (amino acids 1-146) containing the Tudor domain. In the absence of any peptide, no GST-SMN protein accumulated on streptavidin beads (Fig. 5D, lane 1). Importantly, the SmE-N(Me_2_s) peptide, but not the unmodified peptide, readily bound the SMN fragment (Fig. 5D, lanes 2 and 3), indicating that symmetrically dimethylated Arg4 of SmE is recognized by the SMN Tudor domain. **Structural studies on the methylosome-U7 6S complex.** Methylation of SmE by both recombinant and endogenous methylosome prompted us to assemble entirely recombinant complex consisting of U7-specific 6S (U7 6S) intermediate and the methylosome, and to test whether it is amenable for structural studies. The U7 6S intermediate was generated by mixing Lsm10/11 heterodimer, SmE/F/G heterotrimer and pICln, and shown to survive gel filtration chromatography (Fig. 6A), confirming that the hexameric complex is stable. The U7 6S intermediate together with recombinant PRMT5/MEP50 formed a larger complex that eluted as a single peak on gel filtration chromatography (Fig. 6A). Interestingly, gel filtration studies suggest that the methylosome heterooctamer binds only one copy of the U7 6S complex (4:1 molar ratio of PRMT5/MEP50 to U7 6S). The same ratio was observed when a large molar excess of the U7 6S complex was used, suggesting that the methylosome contains only one active binding site for the U7 6S. When applied to the cryo-EM grids, the complex became disrupted, with only the methylosome remaining stable. To stabilize the complex, we used cross-linking with 4 mM BS3 but failed to observe any density for SAM in the EM map. We repeated the crosslinking with 8 mM BS3 in the presence of the SAM analog inhibitor sinefungin (SFG). This approach allowed us to determine a structure of the methylosome at 2.69 Å resolution (Fig. 6B, Table S1), yielding well-defined EM density for the adenosine portion of SFG. The rest of SFG had weaker density and assumed different conformations in the four PRMT5 active sites.

**Fig. 6.**
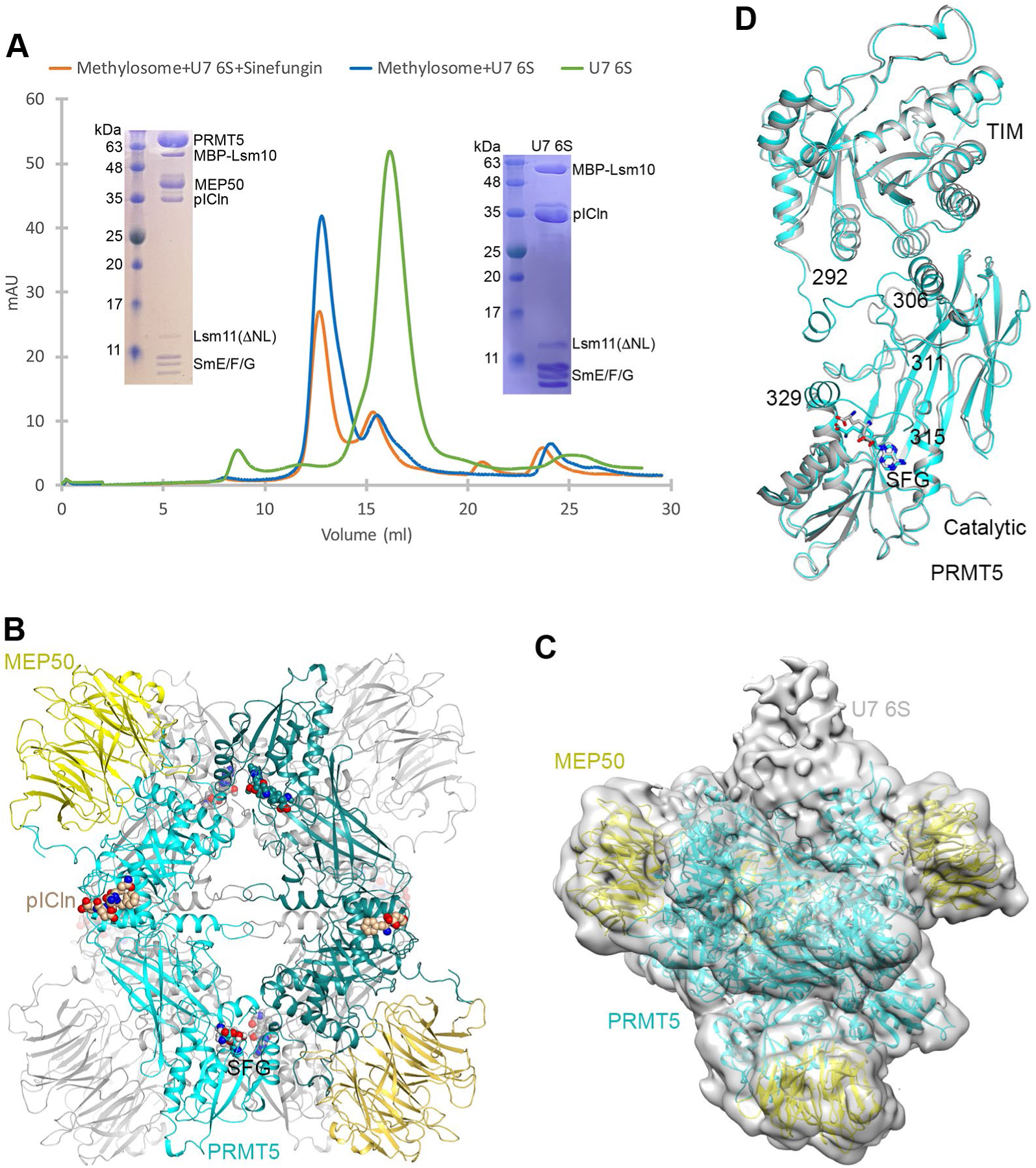
Structural studies of the human methylosome-U7 6S complex. **A.** Gel filtration profiles of Lsm10/11 bound to SmE/F/G and pICln (U7 6S complex, green), methylosome bound to U7 6S (blue), and methylosome bound to U7 6S in the presence of sinefungin (SFG, orange). Inserts are SDS gels of the purified U7 6S sample (right) and methylosome- U7 6S sample (left). Lsm10 is N-terminally fused with MBP, and the N-terminal region and internal loop of Lsm11 are deleted (Lsm11ΔNL). **B.** Schematic drawing of the human methylosome-U7 6S complex. Only the PBM of pICln is included in the atomic model, and the rest of the U7 6S is flexible. **C.** Cryo-EM density for the methylosome-U7 6S complex, low pass filtered to 7 Å resolution. The atomic model for the methylosome is shown. The extra EM density that could correspond to U7 6S is labeled. **D.** Overlay of two PRMT5 molecules with different conformations in the methylosome-U7 6S complex. The PRMT5 molecule in contact with the putative U7 6S EM density is shown in cyan and the other molecule in gray. The disordered segments, 292-306 and 311-315 or 311-329, are labeled. Panel D was produced with Chimera (Goddard et al. 2007).

Weak density was observed associated with one of the PRMT5 molecules in the EM reconstruction (Fig. 6C). The density could not be conclusively interpreted as the U7 6S, although the stoichiometry was consistent with gel filtration studies. The binding of U7 6S to the methylosome is likely highly dynamic. The density binds one PRMT5 molecule and has no contact with MEP50. It is not clear based on the map why there is only one U7 6S binding site in the PRMT5 tetramer.

The structural analysis indicates that the four PRMT5 molecules in the methylosome assume two distinct conformations, with rms distance of 0.4 Å within each pair of the molecules with similar conformation. In contrast, the rms distance between two PRMT5 molecules with different conformation is 0.9 Å, and the differences are mostly located at the interface between the catalytic and TIM barrel domains of PRMT5 (Fig. 6D). This interface is also where the density for U7 6S is located, and residues 292-306 that link the two domains are ordered in the region where U7 6S contacts PRMT5 but disordered in the other PRMT5 molecule. These residues are also disordered in the structure of the methylosome-Lsm10/11 complex, although they are ordered in the complex with the histone H4 peptide (Antonysamy et al. 2012).

In the active site region, part of the segment containing residues 311-325 next to the SFG molecule is disordered, and there is no EM density for a peptide substrate. However, the extra density that could correspond to U7 6S is located near the active site region, making it possible for the N-terminal segment of SmE to reach the PRMT5 active site. This putative U7 6S density is also close to the PBM of pICln, although there is a long linker (80 residues) between the Sm fold of pICln in the U7 6S ring and the PBM at its C terminus.

## DISCUSSION

Spliceosomal snRNPs of the Sm-type (U1, U2, U4, U5, U11, U12 and U4_ATAC_) share the same ring-shaped core domain that surrounds the Sm-binding site in their cognate snRNA component (Will and Luhrmann 2001). The ring consists of seven Sm proteins, D1, D2, D3, B, E, F and G, which are loaded around the Sm-binding site as three pre-formed sub- complexes, SmD1/D2, SmB/D3 and SmE/F/G (Lührmann et al. 1990; Khusial et al. 2005). This assembly is controlled by two large macromolecular entities: the PRMT5 methylosome consisting of three subunits (PRMT5, MEP50, pICln), and the SMN complex consisting of nine known subunits (SMN, Gemin2-8 and Unrip) (Battle et al. 2006a; Neuenkirchen et al. 2008; Li et al. 2014; Gruss et al. 2017). Initially, pICln acts outside the methylosome complex by pre-organizing the Sm subunits into intermediate subcomplexes (Chari et al. 2008; Paknia et al. 2016). In the next step, pICln in conjunction with two remaining methylosome subunits mediate symmetrical dimethylation of arginines in RG-rich C-terminal tails of the Sm proteins D1, B and D3. These modifications serve to recruit the SMN protein, which together with other subunits of the SMN complex controls subsequent steps in the assembly process, including the recognition of a spliceosomal snRNA and its orderly binding to Sm proteins.

U7 snRNP contains a unique Sm ring in which Sm proteins D1 and D2 are replaced by Lsm10 and Lsm11 (Schumperli and Pillai 2004; Sun et al. 2020). These two specific proteins and U7 snRNA are key determinants of the unusual function of U7 snRNP, which plays no recognized role in splicing and instead acts as an RNA-guided endonuclease that cleaves replication-dependent histone pre-mRNAs during their 3’ end processing (Dominski and Tong 2021). The remaining five subunits of the U7 ring are shared with the spliceosomal snRNPs. The Sm-binding site in U7 snRNA partially departs from the consensus established for the spliceosomal snRNAs, promoting the incorporation of Lsm10 and Lsm11 into the ring (Pillai et al. 2003). U7 snRNP is relatively rare in animal cells, being ∼500-fold less abundant than the spliceosomal snRNPs (Smith et al. 1991). Clearly, the assembly of U7 snRNP must be under strict cellular surveillance to prevent binding of the highly abundant SmD1/D2 heterodimer to U7 snRNA, yielding dysfunctional and potentially harmful complexes. Previous *in vitro* and *in vivo* studies established that the assembly of the U7-specific Sm ring on U7 snRNA utilizes at least some components of the PRMT5 and SMN complexes but how exactly they work together to achieve this goal and to distinguish U7 snRNP from the spliceosomal snRNPs is unknown (Schumperli and Pillai 2004; Tisdale et al. 2013; Tisdale et al. 2022). Here, we took a closer look at the role of the PRMT5 methylosome in this process.

### Interaction of the Lsm10/11 heterodimer with the PRMT5 methylosome

In our studies we took advantage of the Lsm10/11 heterodimer that was expressed using baculovirus system and previously used in generating catalytically active semi- and fully recombinant U7 snRNP for structural and functional studies (Bucholc et al. 2020; Sun et al. 2020; Yang et al. 2020). When incubated in cytoplasmic or nuclear extracts from mammalian cells, the heterodimer tightly interacts with endogenous methylosome complex consisting of all three known subunits: PRMT5, MEP50 and pICln. The Lsm10/11 heterodimer also interacts with recombinant methylosome reconstituted from baculovirus-expressed PRTM5/MEP50 heterodimer and bacterially expressed pICln, facilitating structural studies by cryo-EM. In the structure, PRMT5 and MEP50 form a heterooctamer of four tightly bound heterodimers, consistent with previous structural studies of the methylosome by X-ray crystallography and cryo-EM (Antonysamy et al. 2012; Ho et al. 2013; Timm et al. 2018). In the active site of PRMT5, we identified density likely corresponding to residues 71- GRGRGR-76 of Lsm11, with the Arg74 and/or Arg72 side chain located next to the SAM methyl group donor and poised for methylation. This peptide is part of a larger RG cluster located between amino acids 68-79 in Lsm11, RGGGRGRGRARG, that resembles the C-terminal RG clusters found in Sm proteins D1, B and D3. Other parts of the complex were largely disordered, providing no structural information.

By using various mutants of Lsm11 in pull-down assays, we mapped the binding site for the methylosome to Arg74 (primary site) and Arg72 (secondary site), consistent with the structural studies. The methylosome strongly interacts with the Lsm10/11 heterodimer lacking the entire N-terminal region of Lsm11, including the RGGGRGRGRARG cluster. In this case, the interaction entirely depends on the presence of pICln, suggesting the existence of a pICln-mediated contact between the methylosome and the Lsm10/11 heterodimer. Our cryo-EM structure suggests that pICln uses its PRMT5 binding motif (PBM) to stabilize the interaction between Lsm10/11 and the methylosome, consistent with a previous study (Mulvaney et al. 2021). What remains undetectable in the structure is how pICln interacts with the Lsm10/11 heterodimer. Important clues on this interaction come from previous structural studies on the spliceosomal 6S complex consisting of the Sm proteins D1, D2, E, F and G and pICln (Friesen et al. 2001b; Chari et al. 2008; Grimm et al. 2013; Pelz et al. 2015). In this complex, pICln uses β4 strand of its pleckstrin homology domain to directly contact β5 strand of SmD1 in an antiparallel orientation, thus mimicking the interaction mode that oligomerizes Sm proteins (Kambach et al. 1999). The most likely possibility consistent with our binding results is that the same interaction occurs in the U7 6S complex, bringing together Lsm10 (the SmD1 counterpart) and pICln (Paknia et al. 2016).

### Lsm11 is methylated by PRMT5

The presence of the GR cluster of Lsm11 in the active site of PRMT5 strongly suggests that one or more residues of this protein are methylated. *In vitro* methylation assay with ^3^H-labelled SAM and either endogenous or recombinant PRMT5 methylosome confirmed this prediction, demonstrating that Lsm11, but not Lsm10, is a substrate for methylation by the methylosome. Methylation of Lsm11 does not require the pICln subunit and primarily occurs at Arg74, and with lower efficiency at Arg72. Both residues are flanked by glycines, thus exist in an optimal context known to support highly efficient methylation by PRMT5 (Musiani et al. 2019). Simultaneous substitutions of Arg72 and Arg74 with alanines abolished stable interaction of the N-terminal region of Lsm11 with the methylosome, hence preventing Lsm11 methylation. We anticipate that in the presence of Lsm10 that recruits the methylosome via pICln, the double mutant of Lsm11 may undergo methylation at other arginines within the RGGGRGRGRARG peptide and potentially elsewhere in the N-terminal region of Lsm11 (amino acids 1-154). However, the fact that no detectable *in vitro* methylation occurs in the absence of this region (e.g., in Lsm10/11ΔNL) suggests that other parts of Lms11 (and Lsm10) lack suitable methylation sites.

Based on both biochemical and structural studies, we conclude that Lsm11 becomes methylated *in vitro* by PRMT5 within its N-terminal region, contrasting with the Sm proteins D1, B and D3, which are methylated C-terminally. Our results differ from a previous report that failed to identify methylation of Lsm11, likely due to using relatively small amounts of this protein generated *in vitro* (Azzouz et al. 2005).

### Methylation of SmE

Methylation of Lsm11 in U7-specific 6S complex consisting of Lsm10, Lsm11, SmE/F/G heterotrimer and pICln was largely unaffected, but surprisingly an additional protein was methylated in the complex, undetectable in the absence of pICln and/or SmE/F/G. This protein was identified as SmE, with methylation site mapped by mutagenesis to the N-terminal arginine that is followed by a glycine (amino acids 4-5). This RG dipeptide is located within a 15-amino acid unstructured N-terminal extension of SmE that is highly conserved among distant vertebrates, indicative of an important biological function.

Methylation of SmE was readily catalyzed *in vitro* by both recombinant and endogenous methylosome purified from HeLa and mouse cytoplasmic extracts, making it unlikely to be an artifact of PRMT5 overexpression or misfolding. In addition, the same *in vitro* methylation reaction carried out with the spliceosome-specific 6S complex, consisting of SmD1/D2 heterodimer, SmE/F/G heterotrimer and pICln (Grimm et al. 2013) resulted in methylation of only SmD1, consistent with previously reported data (Neuenkirchen et al. 2015). Thus, methylation of SmE specifically depends on the presence of Lsm10 and/or Lsm11 in the complex. Surprisingly, the N-terminal region of Lsm11, the most characteristic feature distinguishing Lsm11 from all other Sm/Lsm subunits and the functional hub of the U7 snRNP, is dispensable for SmE methylation. Presumably, the U7- specific 6S intermediate containing Lsm10 and Lsm11 adopts a different conformation than the spliceosomal 6S complex with SmD1 and SmD2, exposing the N-terminal region of SmE for the enzymatic activity of PRMT5.

To gain additional information on how PRMT5 methyltransferase accesses the N-terminal arginine in SmE for methylation in the U7-specific 6S complex, we carried out a set of studies using cryo-EM. These studies were largely unsuccessful due to instability and/or flexibility of the complex. Nevertheless, the available densities are consistent with the predicted juxtaposition of the catalytic center of PRMT5 with the N-terminal part of SmE. Interestingly, both gel filtration chromatography and cryo-EM models indicate that of the four copies of the PRMT5/MEP50 heterodimer in the methylosome, only one is accessible for the U7 6S complex. The reason for this sub-stoichiometric binding is unknown. Perhaps, the U7 6S complex bound to one copy of the PRMT5/MEP50 heterooctamer imposes structural changes on the remaining three copies, preventing their interaction with the complex. In support of this interpretation, our structural studies revealed some conformational changes in PRMT5.

### Potential roles of Lsm11 and SmE methylation in U7 snRNP biogenesis

The most important question for future studies is whether methylation of Lsm11 and SmE occurs *in vivo*. Our initial attempts with endogenous U7 snRNP partially purified from mouse and human nuclear extracts failed to detect the presence of sDMA modification on either of the two proteins by Western blotting using sDMA-specific antibodies Y12 and SYM11. The same antibodies detected Sm protein B and D3 in the purified preparation of U7 snRNP (data not shown). However, compared to the multiple RG repeats in SmB and SmD3, the RG-rich track in Lsm11 is relatively short, and the methylation site in SmE contains only a single arginine, likely forming epitopes too weak to be recognized by either antibody. Due to the very limiting nature of endogenous U7 snRNP, our efforts to use mass spectrometry to determine the methylation pattern generated on Lsm11 and SmE *in vivo* were so far unsuccessful.

In conclusion, our *in vitro* results suggest that the assembly of the U7 snRNP involves specific modification of its two subunits by the PRMT5 methylosome: Lsm11 and SmE. Both subunits are methylated within short, N-terminal motifs, contrasting with the much longer and C-terminally located methylation targets in the Sm proteins D1, B and D3. Symmetrically dimethylated arginines interact with the Tudor domain of the SMN protein (Friesen et al. 2001a; Selenko et al. 2001; Cote and Richard 2005; Tripsianes et al. 2011), facilitating the transfer of the Sm proteins to the SMN complex and efficient and faithful assembly of the Sm ring on the spliceosomal snRNAs (Gruss et al. 2017). It can be assumed that methylation of Sm proteins in the U7-specific ring serves the same purpose, with the unique pattern of methylated arginines on Lsm11 and SmE likely playing a key role in distinguishing the U7 snRNP assembly pathway from the much more general and ubiquitous assembly pathway for the spliceosomal snRNPs.

## MATERIALS and METHODS

### Antibodies

Rabbit serum containing antibodies against a dimer of Lsm10-MBP and full length Lsm11 was generated by Pacific Immunology (Ramona, California). The following commercial antibodies were used in this study: SmB 12F5 (Sigma), SmD1 RB17341 (MyBioSource), SmD3 A303-954A (Bethyl), SmE ab229557 (Abcam), Y12 MA-1-90490 (Thermo), pICln A304-521A (Bethyl), MEP50/Mep50 A301-562A (Bethyl), PRMT5 A300-850A (Bethyl), MBP 66003-1-Ig (Proteintech), Symmetric Dimethyl-Arginine SYM11 (Millipore).

### Cell culture and preparation of cytoplasmic and nuclear extracts

HeLa and mouse myeloma cells were grown by Cell Culture Company (Minneapolis, MN). HeLa cells were shipped on dry ice as a frozen cell pellet and used after thawing. Myeloma cells were shipped as a concentrated cell suspension on wet ice and used immediately upon arrival. Nuclear extracts from HeLa and mouse myeloma cells were prepared, as described (Dominski et al. 1995; Skrajna et al. 2018; Sun et al. 2021). The crude cytoplasmic fractions generated after collecting HeLa or mouse myeloma nuclei were used to prepare cytoplasmic extracts, as described (Mayeda and Krainer 1999), with the ultracentrifugation and dialysis steps being omitted.

### Purification of pICln

Full-length mouse pICln was cloned into the pET28a (Novagen) vector with a 6xHis tag added to the N terminus and expressed in *E. coli* Rosetta strain (Novagen). The cell pellet was resuspended and lysed by sonication on ice in 100 mL of lysis buffer containing 25 mM Tris (pH 7.5), 300 mM NaCl, and 5% (v/v) glycerol. The supernatant was incubated with nickel beads for 1 h at 4 ℃. The beads were washed 2 times with 50 bed volumes of wash buffer containing 25 mM Tris (pH 7.5), 300 mM NaCl, and 20 mM imidazole. The protein was eluted with 5 mL of elution buffer containing 25 mM Tris (pH 7.5), 300 mM NaCl, and 250 mM imidazole. Eluate was diluted 3X with 25 mM Tris (pH 7.5) and was purified by chromatography using a HiTrap MonoQ column (Cytiva) with a salt gradient. Buffer A contained 25 mM Tris (pH 7.5) and 100 mM NaCl, and Buffer B contained 25 mM Tris (pH 7.5) and 1 M NaCl. Fractions of interest were concentrated to ∼1.1-1.7 mg/mL with 5% (v/v) glycerol, flash frozen using liquid nitrogen, and stored at –80 ℃.

### Purification of the PRMT5/MEP50 complex

cDNAs encoding full-length human PRMT5 and MEP50 were cloned into pFL vector and co-expressed in insect cells. One liter of High5 cells (1.8x10^6^ cells/mL) was infected with 15 mL of PRMT5/MEP50 P2 virus with the His tag on PRMT5. For purification, the cell pellet was resuspended and lysed by sonication on ice in 100 mL of lysis buffer containing 25 mM Tris (pH 7.5), 300 mM NaCl, 5% (v/v) glycerol, and one protease inhibitor cocktail tablet (Sigma). The cell lysate was centrifuged at 13,000 rpm for 45 mins at 4℃. The supernatant was incubated with nickel beads for 1 h at 4 ℃. The beads were washed 2 times with 50 bed volumes of wash buffer and eluted with 5 mL of elution buffer, as described for pICln. The eluate was purified by gel filtration using a Superdex 200 Increase 10/300 GL column (Cytiva) in buffer containing 25 mM Tris (pH 7.5), 150 mM NaCl, 2 mM DTT, and 10% (v/v) glycerol. Fractions of interest were concentrated to ∼1.1 mg/mL with 5% (v/v) glycerol, flash frozen using liquid nitrogen, and stored at –80 ℃.

### Purification of the Lsm10/11 heterodimer

The expression and purification of human Lsm11 lacking residues 211-322, and full length Lsm10 was carried out as described previously (Bucholc et al. 2020; Sun et al. 2020; Sun et al. 2021). Briefly, Lsm11 and Lsm10 were cloned into pFL vector with maltose-binding protein added to the N terminus of Lsm10. One liter of High5 cells (1.8x10^6^ cells/mL) was infected with 15 mL of Lsm10/11 P2 virus. The cell pellet was resuspended and lysed by sonication on ice in 100 mL buffer containing 25 mM Tris (pH 7.5), 500 mM NaCl, 5% (v/v) glycerol, and one protease inhibitor cocktail tablet (Sigma). The cell lysate was centrifuged at 13,000 rpm for 45 mins at 4 ℃. The supernatant was incubated with nickel beads for 1 h at 4℃. The beads were washed 2 times with 50 bed volumes of wash buffer (25 mM Tris (pH 7.5), 500 mM NaCl, and 40 mM imidazole) and eluted with 5 mL of elution buffer containing 25 mM Tris (pH 7.5), 500 mM NaCl, 500 mM imidazole, and 5% (v/v) glycerol. The eluate was diluted 2.5X with 25 mM HEPES (pH 7.5) and 5 mM DTT. The complex was then purified by chromatography using a HiTrap Heparin column (Cytiva) with a salt gradient. Buffer A contained 20 mM Hepes (pH 7.5) and 5 mM DTT, and Buffer B contained 20 mM Hepes (pH 7.5), 5 mM DTT, and 1 M NaCl. Fractions of interest were concentrated to ∼1 mg/mL with 5% (v/v) glycerol, flash frozen using liquid nitrogen, and stored at –80 ℃.

### Purification of endogenous methylosome bound to Lsm10-MBP/Lsm11 heterodimer or Lsm11 alone

MBP-Lsm10/Lsm11 heterodimer or various N-terminal fragments of Lsm11 fused to MBP (depending on experiment, 25-100 pmol each) were incubated for 1 hour on ice with 750 µl of cytoplasmic or nuclear extracts from mouse myeloma or HeLa cells. The samples were supplemented with 5 µl of a rabbit serum generated against -MBP Lsm10/Lsm11 heterodimer, rotated 60 min in cold room and spun down in a microcentrifuge (10 min at 10,000 x *g*) to remove potential precipitates. The supernatants were loaded over 30 µl of protein A plus beads (Pierce), rotated in cold room for 75 min and gently spun to collect beads and the bound complexes. The beads were washed for 60 min with buffers matching those present in the cytoplasmic or nuclear extracts, moved to new tubes for an additional 30 min rotation and resuspend in SDS sample buffer. A fraction of each sample was separated in a 4-12% SDS/polyacrylamide gel and immunoprecipitated proteins were visualized by silver staining and identified by mass spectrometry in the Laboratory of Mass Spectrometry at the Institute of Biochemistry and Biophysics (Warsaw Poland), as previously described (Yang et al. 2013; Skrajna et al. 2018). Protein identities were confirmed by Western blotting using specific antibodies. In some experiments, instead of using anti-Lsm10/11 serum, proteins bound to the Lsm10 /Lsm11 heterodimer were directly collected on amylose beads. In this approach, extracts mixed with MBP-Lsm10/Lsm11 heterodimer were rotated 60 min in cold room, spun down in a microcentrifuge, as above, and rotated with 30 µl of amylose beads. The remaining steps were the same as during the immunoprecipitation protocol.

### *In vitro* methylation by recombinant methylosome

Recombinant methylosome consisting of baculovirus-expressed PRMT5/MEP50 and bacterially expressed pICln (1 pmol each) was mixed in 20 µl of the cytoplasmic buffer with 2- to 10-fold excess of recombinant Sm proteins and each sample was supplemented with 2 µCi of ^3^H-labeled SAM (PerkinElmer) for overnight methylation at 32 °C. Each sample was mixed with equal volume of 2x SDS sample buffer, resolved in 15% or 4-12% SDS/polyacrylamide gels. Separated proteins were visualized by Coomassie Blue staining and gel images captured. The stained gels were incubated for 30 min with Amplify solution (GE Healthcare), dried and used for 12 to 72-hour fluorography at -80 °C. Methylated proteins were identified by aligning fluorograms with stained gel images.

### *In vitro* methylation by endogenous methylosome

MBP-Lsm10/Lsm11 heterodimer (100 pmol) either alone or together with 300 pmol of SmE/F/G heterotrimer was rotated for 60 min in cold room with Hela or mouse myeloma cytoplasmic extracts (750 µl) and spun down at 10,000 x *g* for 10 min. The heterodimer bound to endogenous methylosome was purified on amylose beads, as described above. After extensive washing, the beads were suspended in 25 µl of cytoplasmic buffer containing 2 µCi of ^3^H-labeled SAM (PerkinElmer) and incubated overnight at 32 °C without shaking. Following incubation, the supernatant was removed, and the beads were resuspended in 25 µl of SDS sample buffer. Proteins collected on the beads were resolved in 15% polyacrylamide or 4-12% SDS/polyacrylamide gels and stained with Coomassie Blue. Protein composition of bound complexes was additionally determined by silver staining and/or Western blotting. Arginine methylation was detected by fluorography, as described above. In some experiments, this approach was also used to immobilize recombinant methylosome to conduct the methylation assay on amylose beads rather than in solution.

### EM studies of the methylosome bound to the Lsm10/11 heterodimer

Purified PRMT5/MEP50, Lsm10/11 and pICln were mixed on ice (1:1:1.2 molar ratio) and incubated for 1 h. The resultant complex was purified by size exclusion chromatography using Superose 6 column (Cytiva) in a buffer containing 25 mM Tris (pH 7.5), 150 mM NaCl, and 2 mM DTT. Fractions of interest were concentrated to ∼0.5 mg/mL, flash frozen with liquid nitrogen, and stored at –80 ℃. Cryo-EM grids were prepared by applying 3.5 μL of protein sample at a concentration of 0.25 mg/ml to one side of a Quantifoil 400 mesh 1.2/1.3 gold grid with graphene oxide support film (Quantifoil). After 30 s, the grid was blotted for 1.5 s on the other side under 99% humidity and at 20 °C using EM GP2 plunge freezer (Leica) and immediately plunged into liquid ethane. 4,601 image stacks were collected on a Titan Krios electron microscope at the Columbia University Cryo-Electron Microscopy Center, equipped with a K3 direct electron detector (Gatan) at 300 kV with a total dose of 58.2 e Å^−2^ subdivided into 50 frames in 2.5 s exposure using Leginon. The images were recorded at a nominal magnification of 105,000× and a calibrated pixel size of 0.83 Å, with a defocus range from –0.8 to −2.2 μm. Image stacks were motion-corrected and dose-weighted using RELION 3.1 (Zivanov et al. 2018). The patch CTF parameters were determined with cryoSPARC (Punjani et al. 2017). First, 337,438 particles were auto-picked from 500 images and were used to generate six 3D initial models by *ab initio* reconstruction. The model with recognizable features by visual inspection was chosen for creating templates for template picking. 1,814,708 particles were picked from 4,601 micrographs by template picking. After two rounds of heterogeneous refinement, 754,047 particles were imported to RELION for CTF refinement and Bayesian polishing. The polished particles were then imported back to cryoSPARC for a homogeneous refinement, yielding a map at 2.86 Å resolution. C1 symmetry was used throughout the reconstruction. The crystal structure of PRMT5/MEP50 bound to the PRMT5 binding motif (PBM) peptide of pICln (PDB code 6V0O) was fitted as a rigid body into the EM map using Chimera (Pettersen et al. 2004). The atomic model of the Lsm11 peptide (residues 71-76) was built manually into the cryo-EM density with Coot (Emsley and Cowtan 2004). The atomic model was improved by real-space refinement with the program PHENIX (Liebschner et al. 2019). The cryo-EM information is summarized in Table S1.

### Purification and EM studies of the methylosome/U7 6S complex

Human SmE/F/G heterotrimer and human MBP-Lsm10/11ΔNL heterodimer were expressed in *E. coli* and insect cells, respectively and purified as described earlier (Bucholc et al. 2020; Sun et al. 2020; Sun et al. 2021). Lsm10 carried an N-terminal maltose binding protein (MBP) and a 6xHis tag. In ΔNL Lsm11, the N-terminal segment (residues 1-152) and the internal loop (residues 211-322) were removed. Purified SmE/F/G heterotrimer, MBP-Lsm10/ΔNL heterodimer and mouse pICln were mixed (molar ratio 1.2:1:1.2) in a buffer containing 20 mM Tris (pH 8.1), 100 mM NaCl and 2 mM DTT and incubated on ice for 1 h. The assembled U7 6S complex was purified with a Superose 6 (Cytiva) gel filtration column using the same buffer. Purified PRMT5/MEP50 and U7 6S were combined (molar ratio 4:8 or 4:2) in a buffer containing 20 mM Hepes (pH 7.7), 100 mM NaCl and 2 mM DTT and incubated on ice for 1 h. The assembled methylosome-U7 6S complex was purified with a Superose 6 (Cytiva) gel filtration column. To stabilize the methylosome-U7 6S complex, the sample was cross-linked on ice for 2 h using 4 mM BS3 allowing cryo-EM reconstruction at 3.7 Å resolution. In some experiments, we used the SAM analog sinefungin at 0.5 μM and 1.5 h cross-linking with 8 mM BS3. The cryo grids were prepared following the protocols described above, and the cryo-EM data were collected using a Titan Krios microscope at the New York Structural Biology Center. The EM data processing followed the protocols described above, and the statistics are summarized in Table S1. Weak, putative EM density for U7 6S was observed after heterogeneous refinement in cryoSPARC, while the density became much weaker after non-uniform or homogeneous refinement. Various attempts to improve the quality of the density were unsuccessful.

## SUPPLEMENTAL MATERIAL

Supplemental material is available for this article.

## ACKNOWLEDGEMENTS

We thank J. Oledzki and A. Fabijanska (Laboratory of Mass Spectrometry, Institute of Biochemistry and Biophysics, Polish Academy of Sciences) for mass spectrometry analysis, and K. Krajewski (High-Throughput Peptide Synthesis and Array Core Facility, UNC, Chapel Hill) for peptide synthesis. We also thank L. Pellizzoni (Columbia University) for the gift of a monoclonal anti-SmB antibody. This research was funded by the National Institutes of Health (NIH) grants GM 29832 (Z.D.) and R35GM118093 (L.T.).

